# Multiplexed Amplicon Sequencing Reveals High Sequence Diversity of Antibiotic Resistance Genes in Québec Sewers

**DOI:** 10.1101/2023.03.06.531290

**Authors:** Claire Gibson, Susanne A. Kraemer, Natalia Klimova, Laura Vanderweyen, Nouha Klai, Emmanuel Díaz Mendoza, Bing Guo, David Walsh, Dominic Frigon

## Abstract

The United Nations Environment Assembly (UNEA-3) have recognised the importance of the environment in the development, spread and transmission of antimicrobial resistance (AMR) to humans and animals. Such recognition calls for wider surveillance of antimicrobial resistance genes (ARG) in wastewater and other environmental reservoirs. For ARG surveillance to be valuable to regulators, it must enable source tracking and risk assessment. Adequate surveillance also requires the processing of a large number of samples at a relatively low cost, and a low detection limit to allow quantification of the riskiest ARGs. However, current methods for tracking ARGs have various limitations. The current study presents a multiplexed targeted amplicon sequencing approach for the detection of sequence variants of ARGs in environmental samples. To demonstrate the application of this technique, wastewater samples collected from the inlet to 16 treatment plants located along a 440-km transect of the St-Lawrence river in the province of Quebec (Canada) were analysed. Among the ARGs examined, between 3 and 45 nucleic acid sequence variants were detected demonstrating the high sequence diversity that occurs within genes originating from a single sample type and the information that is missed using traditional techniques. Using the PLSDB and Comprehensive Antibiotic Resistance Database (CARD), the risk of ARG sequence variants was inferred based upon their reported mobility and detection in pathogens. Results suggest that sequence variants within a single ARG class present different risks to public health. In the future, targeted amplicon sequencing could be a valuable tool in environmental studies for both risk assessment purposes and in AMR source tracking.

## 1.0 Introduction

Antimicrobial resistance (AMR) is widely recognised as one of the greatest threats to modern medicine of the twenty first century. Traditional AMR monitoring strategies focused on areas with high antibiotic usage such as clinical and agricultural settings. More recently, the importance of the environment in the development, spread and transmission of AMR to humans and animals has emerged (1,2). One of the most notable examples is the *bla*NDM gene which was first identified in 2008 (3), and was demonstrated to disseminate in the environment through wastewaters (4). Such recognition calls for wider surveillance of antimicrobial resistance genes (ARG) in wastewater and other environmental reservoirs.

Studies are increasingly recognising the value of monitoring ARG sequence diversity for source tracking purposes. Machine learning classification has been used to profile the sequences of ARGs from different environments and train models to determine potential source types (5,6). Both common and unique ARGs were identified across ecotypes highlighting the potential value of obtaining data on ARG sequence diversity (6). To improve the performance of source tracking models and allow the tracking of resistome development in specific areas, it has been suggested that regional studies of ARGs are required. However, this can be costly and the data handling cumbersome. Zhang et al., 2021 studied amino acid sequence variants of ARGs in different habitats including feces, WWTPs and environmental samples from non-anthropogenically impacted areas. Assessment of ARG risk revealed that ARGs in the same gene family can pose substantially different risks due to differing mobility, hosts and ecological distribution (7). Taken together, these studies demonstrate the valuable information that can be gained and potential applications of ARG sequence variant monitoring. Yet, to date these approaches are limited due to the high cost per sample and the lengthy analyses required.

Current methods for the detection of ARGs in environmental samples have various limitations. Shotgun metagenomics can detect a large number of ARGs and mobile genetic elements, but the detection limit is often poor (10^-3^ copy/genome) and the analysis requires high levels of training. This method can also be costly, often meaning that fewer samples are analysed, which in turn limits the construction of large datasets which are required to understand the connectivity between environmental reservoirs. Quantitative PCR techniques have a higher detection limit (10^-5^ to 10^-7^ copy/genome), but further characterisation of sequence diversity in genes can be challenging.

Here, we introduce a multiplex targeted amplicon sequencing approach for the detection of sequence diversity of ARGs in environmental samples. A selection of antibiotic and metal resistance genes were amplified in a multiplex PCR reaction, barcoded to enable sample pooling and sequenced on the Illumina Miseq platform. This method is similar to that developed independently by Smith et al., 2022 (8) and enables relatively fast and low-cost analysis of sequence diversity in multiple ARGs and samples in a single sequencing run. This protocol can also be easily adapted to target different genes depending on the users’ aims, enabling versatility and exhaustive performance calibration in the analysis of the targeted resistome.

To demonstrate the use of multiplex targeted amplicon sequencing, we surveyed influent wastewater from 16 wastewater treatment plants located in the province of Quebec, Canada. Wastewater has been demonstrated to contain a plethora of ARGs and antimicrobial resistant bacteria (ARBs). Previous studies have identified genetic diversity in ARGs based on the sample type and location (6). ARGs in influent wastewater are thought to originate from numerous sources including domestic and clinical waste, industrial discharges, urban run-off and growth selection in sewer biofilms. Consequently, it was hypothesised that ARG sequence diversity would be observed among the sewer samples collected using our novel amplicon sequencing approach. Influent wastewater offers many potential pathways for the environmental dissemination of antibiotic resistance, through combined sewer overflow whereby untreated wastewater is discharged directly into the environment or incomplete removal during the wastewater treatment process. Thus using this data, we explore the risk of different ARG sequence variants and the potential applications of this tool.

## 2.0 Materials and Methods

### 2.1 Sample Collection

Grab samples were collected from the influent wastewater inlet at 16 Québec wastewater treatment plants between June and August 2018. Wastewater samples were transported to McGill University on ice and processed within 24 hours of collection. Samples were centrifuged in 2 mL microcentrifuge tubes at 12,000 ×g for 5 min., the supernatant was then removed, and centrifuged solids were stored at −80 °C until further analysis. Wastewater treatment plant metadata for the sampling period was obtained through the Québec Data Partnership for wastewater treatment plants (9). Samples obtained were assigned a sample ID as reported in Table S1, ordered according to the longitude of the sampling location.

### 2.2 Mapping of Sampling Area

Maps were produced using QGIS software and georeferenced in WSG 84 EPSG:4326. The co-ordinates of the samples WWTP influents were retrieved from publicly available data from the Government of Quebec (10). The co-ordinates of ambulatory health care services, hospitals, and nursing and residential care facilities were retrieved from the open database of health care facilities available on the Government of Canada open database of healthcare facilities (11) and imported to QGIS as a CSV file. Land cover data from 2015 was retrieved from the Government of Canada Website (12), the TIFF file was imported to QGIS and projected from EPSG 3978 NAD83/ Canada Atlas Lambert. The colour scheme for each land category was as presented in the 2015 North American Land cover map (13) and according to the colour scheme outlined in Supplementary Table S2. Cattle numbers were retrieved from the 2016 agricultural census data (14). The TIFF file was imported to QGIS and projected from EPSG:3347 – NAD83/ Statistics Canada Lambert. The census consolidated subdivision level was used with a graduated colour ramp of four quantile (equal count) colour classes.

### 2.3 Microbial Community Analysis

DNA was extracted from stored biomass samples using DNeasy PowerSoil Kit (Qiagen, Germantown, MD, USA). PCR of the 16S rRNA gene V4 region was conducted using the 515F and 806R modified Caporaso primers (15,16). The PCR conditions were as follows: 94 °C for 3 mins followed by 35 cycles of 94 °C for 45 secs, 50 °C for 60 secs, 72 °C for 90 secs. 72 °C for 10 mins and a final hold at 4 °C. After amplification of the 16S rRNA gene, amplicons were barcoded in a second PCR reaction with the following reaction conditions: initial denaturation at 94 °C for 3 minutes, followed by 15 cycles of 94 °C for 30 secs, 59 °C for 20 secs, 68 °C for 45 secs and a final elongation at 68 *°C* for 5 mins. Amplicons were sequenced on the Illumina MiSeq PE250 platform at Génome Québec Innovation Centre (Montréal, QC, Canada). The number of sequences obtained per sample are reported in Table S1.

### 2.4 Multiplexed Amplicon Sequencing

Target Amplicon Sequencing was performed using a custom built accel-amplicon™ Panel designed in collaboration with Swift Biosciences (now Integrated DNA Technologies). This method involved performing a highly multiplexed PCR reaction followed by barcoding and sequencing. The panel used included a total of 159 primer pair combinations targeting 114 ARGs (Table S3). Primers were designed to produce amplicons of similar length with an average of 275 base pairs to avoid preferential amplification of shorter fragments.

Samples were prepared according to the manufacturer’s instructions. An input DNA concentration of 25 ng was used. Magnetic clean-up steps were conducted using SPRI Select Beads (Beckman Coulter, IN, USA). The reaction conditions for the multiplexing PCR were as follows: 98°C for 30s followed by 4 cycles of 98°C for 10s, 63°C for 5 mins, 65°C for 1 min, then 22 cycles of 98°C for 10s and 64°C 1 min, and terminated by a final elongation step of 65°C for 1 min and hold at 4°C. After multiplexing, barcoding and clean-up steps were completed as outlined in the accel-amplicon protocol. Samples were sequenced on the Illumina MiSeq PE250 platform at Génome Québec Innovation Centre (Montréal, QC, Canada).

### 2.5 Bioinformatics

16S rRNA amplicon sequencing data was processed using Qiime2 software (17) and R Software with ‘vegan’ packages (18). Raw sequences were quality filtered using DADA2 (19) and ASV tables extracted from QIIME2 after quality filtering for further analysis in R. Taxonomy was assigned using the MiDAS 2.0 reference database (20) and tabulated. Principal coordinate analysis was performed using R ‘vegan’ package (18) and Bray-Curtis Dissimilarity. A heatmap of the most abundant genera was generated using the function ‘heatmap.2’ in ‘gplots’ package in R (21).

Sequenced Accel-Amplicon samples were obtained from Genome Quebec and forward and reverse reads merged using pear v.0.9.11. Subsequently, the merged reads were assigned to a specific gene product based on their beginning and end (representing the forward and reverse primer sequence used) using a custom python script (available upon request) and all reads converted to fasta format. Reads may have multiple assignments (resulting from degenerate primer sequences). In these cases, we kept the first assignment for each duplicated read. Reads were sorted by length and deduplicated using vsearch v.2.13.3 (22). Unique sequences detected for each target gene may represent real sequence variants or result from PCR and sequencing errors. To filter out such errors, we assumed that the distribution of variant frequencies under an assumption of pure error would follow a Poisson distribution centred around the mean of all counts, with many sequence variants only having one or two reads (which might have been seeded by cross-contamination or misassignments of reads during sequencing). For each sample and gene, we tested whether we could detect sequence variants whose frequencies represented outliers to a Poisson distribution (after correction for multiple testing using the Bonferroni method), representing higher abundance, real sequence variants after PCR. If such sequence variants were detected, we removed low count “noise” sequence variants from the sample by finding the minimum of the distribution function and cutting all sequence variants below the minimum. A custom R script for sequence variant filtering is available upon request. For products in which more than a single variant was retained after filtering, OTU tables were constructed using the clusterfast method in vsearch with an id of 1. To ensure that the sequence variants represented correct products, we used tblastx to compare all sequence variants to a custom database of ARGs based on a dereplicated version of the CARD database with a minimum id of 70 % (blast v.2.12.0). Sequence variants that did not have hits to their respective target gene at this id level were removed.

### 2.6 Multiple Sequence Alignment of ARG Sequences

For multiple sequence alignment, ARG sequences were obtained from NCBI Blast based upon 100 % query coverage and 100 % identity with the ARG sequence variants observed. Results were filtered to include only entries with the gene name included in the description (*bla*TEM or *bla*OXA) and complete CDS only. Sequences were aligned using MAFFT and the fast Fourier transform method (FFT-NS-2) (23) for multiple sequence alignment. Phylogenetic trees were generated using the average linkage (UPGMA) method.

### 2.7 Inverse PCR

Inverse PCR was used as described by Pärnänen et al., 2016 (24) to analyse the flanking regions surrounding the *bla*OXA ARG. Briefly, the unknown flanking regions were digested using restriction enzymes and then ligated together. Clones were then sequenced using Sanger sequencing at Genome Quebec, Montreal. The obtained forward and reverse sequences were analysed using NCBI BLAST (25).

### 2.8 Statistical Analysis

Redundancy analysis (RDA) was used to explain the variation in ARG sequence variants (response variables) due to the explanatory variables considered (influent characteristics, land use and geographical location) (10–12,14). Geographical data (longitude and latitude) was transformed into distance based eigen vector maps (DbMEM) and used as an explanatory variable in the RDA. DbMEM variables were constructed using function dbmem() of package ‘adespatial’ in R (26). The function give.thresh() of package ‘adespatial’ in R was used to select a matrix truncation value of 0.825 to retain only distances among close neighbours. RDA of Hellinger-transformed sequence data for each gene was computed using the ‘rda’ function in R ‘vegan’ package (18). To account for RDA bias which varies based on the number of explanatory variables included, the R^2^ values obtained were adjusted using Ezekiel’s formula. The adjusted R^2^ obtained measured the unbiased explained variation. An unbiased amount of explained variation close to 0 indicates that the explanatory variable did not explain more of the variation in the response variable than random deviates would. The significance of the R^2^ values was tested using function anova.cca in the R ‘vegan’ package (18).

## 3.0 Results and Discussion

### 3.1 Microbial Community of Sewer Samples

In this study, sixteen wastewater samples were analysed which were collected at the entrance of WWTPs distributed along the Ottawa and St. Lawrence rivers between Gatineau and Quebec City (Table S4). The microbial community composition in the wastewater samples was assessed using 16S rRNA gene sequencing. Principle coordinate analysis revealed clustering of the microbial communities (Figure 1a), which did not significantly correlate with operational parameters such as flow and pipe design (all gravity sewer versus presence of pressured lines) or the geographical proximity of samples (Table S4) as assessed using a permutation test of environmental variables (Table S5). Analysis of the overall top 20 most abundant genera in the influent wastewater samples revealed *Acinetobacter* to be dominant members of the microbial community in Clusters 1 and 2 (Figure 1b). This is consistent with previous studies which identify *Acinetobacter* as dominant members of the wastewater microbial community worldwide (27–29). Although many species are non-pathogenic, others such as *Acinetobacter baumannii,* exhibits high levels of AMR to many antibiotics and are a major cause of antimicrobial resistant infection worldwide (30). Taxa from the family *Comamonadaceae,* which are phenotypically diverse and commonly reported in soil and water environments (31), were also dominant in all influent wastewater samples. Another dominant member of the microbial community included *Trichococcus,* a pathogen commonly reported in influent wastewater (32,33), which has been linked to sludge bulking problems in WWTPs (34).

**Figure 1.**
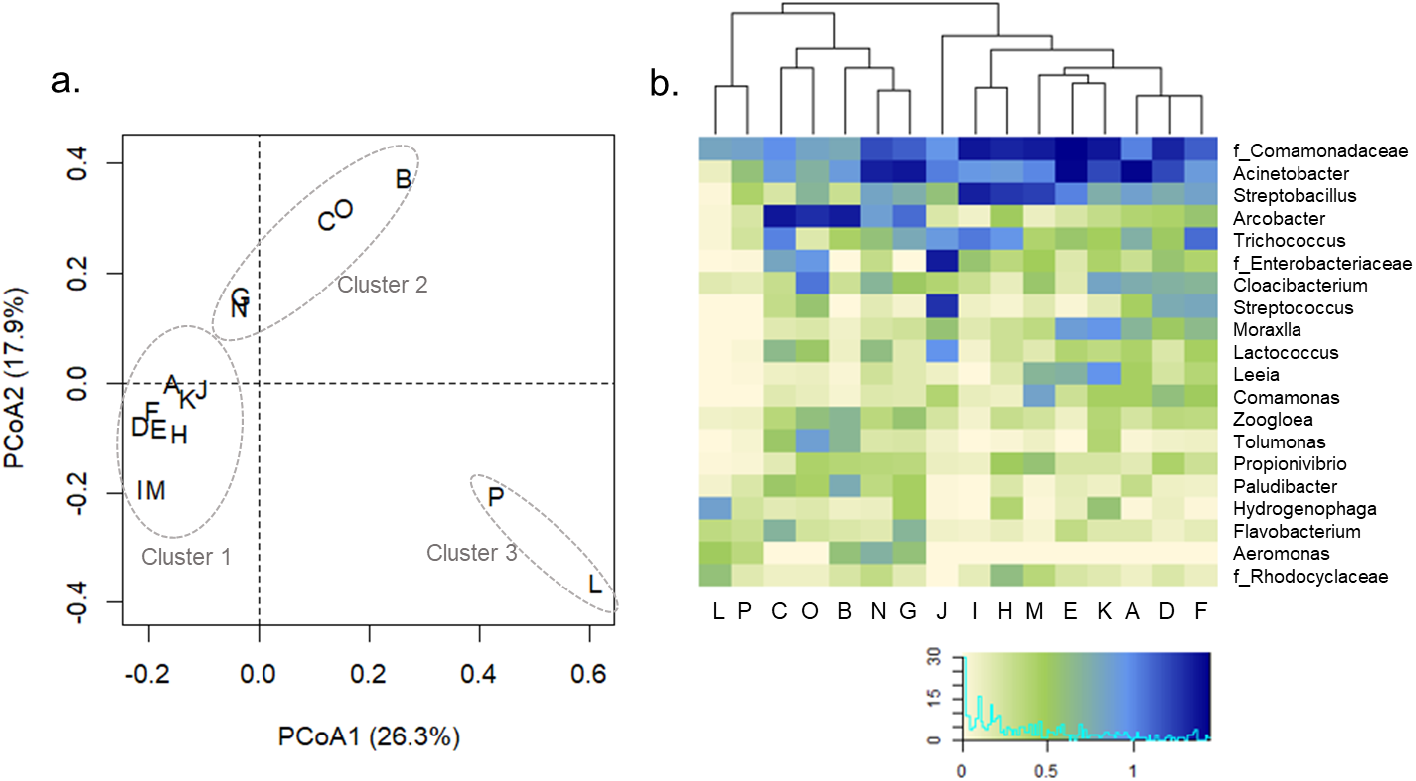
Bacterial community analysis based on amplicon sequencing of the V4 region of the 16S rRNA gene of wastewater samples collected from the inlet to 16 WWTPs in Quebec, Canada. a) Principal coordinate analysis with Bray-Curtis distance b) Heatmap of top 20 most abundant overall genera

Cluster 2 communities (sampling sites: G, N, C, O and B) featured a higher relative abundance of *Arcobacter* than in the communities of Clusters 1 and 3. *Arcobacter* are frequently observed in high abundance in influent wastewater samples (35). *Arcobacter* have been isolated from numerous sources including humans, animals and wastewater and are considered emerging pathogens which have been associated with both human and animal disease (36). Raw wastewater samples have been identified as a potential source of *Arcobacter* infection (37). Finally, Cluster 3 (sampling sites: P and L) were found to harbour communities containing lower relative abundances of *Acinetobacter* and *Arcobacter*. Their distinct microbial communities from the other samples was dominated by *Aeromonas*, which is widespread in the environment and causes a wide range of diseases in humans and animals (38). AMR strains of *Aeromonas* have been demonstrated to persist after the wastewater treatment process and in receiving surface waters (38). Considering the level of AMR in influent wastewater, and the high abundance of important potential pathogens such as *Acinetobacter* and *Arcobacter,* increased surveillance of these environments would be beneficial to manage the associated risk and prevent further dissemination of AMR.

### 3.2 High Sequence Diversity Observed in Sewer ARGs

Using targeted amplicon sequencing, a total of 60 ARGs and metal resistance genes (MRGs) were detected in the sewer samples, 16 of which displayed sequence diversity (Table 1). Of the sixteen genes, between 3 and 45 different sequence variants were observed indicating high genetic diversity in sewer ARGs. Considering the stringent Poisson distribution-based filters used to identify real sequence variants from sequencing errors, it is plausible that the ARG sequence diversity may have been underestimated. The high prevalence of ARG sequence diversity from a single sample type demonstrates the information that is missed when using traditional detection methods such as quantitative PCR.

**Table 1:** Genes detected in sewer samples with sequence diversity and partitioning of variance

| Gene | Resistance Mechanism | # Sewers Detected<sup>a</sup> | # Variants Observed | Unbiased Variation Explained |  |  |  |
| --- | --- | --- | --- | --- | --- | --- | --- |
|  |  |  |  | Influent Factors<sup>b</sup> | Land Factors<sup>c</sup> | Geographical Factors<sup>d</sup> | Microbial Community<sup>e</sup> |
| *arsA* | Metal resistance | 11 | 16 | 0.00 | 0.12 | −0.02 | 0.07 |
| *blaACT* | Beta-lactamase | 13 | 45 | 0.07 | −0.07 | 0.01 | 0.03 |
| *blaOXA* | Beta-lactamase | 16 | 4 | 0.58 | −0.21 | −0.13 | 0.07 |
| *blaSHV* | Beta-lactamase | 9 | 9 | 0.09 | −0.03 | −0.06 | 0.08 |
| *blaTEM* | Beta-lactamase | 16 | 3 | 0.20• | −0.07 | 0.07 | −0.15 |
| *copA* | Metal resistance | 11 | 16 | 0.08 | 0.18• | −0.04 | 0.06 |
| *mdtF* | Efflux pump | 7 | 7 | −0.03 | −0.13 | 0.01 | −0.07 |
| *mdtG* | Efflux pump | 5 | 3 | 0.30* | 0.13 | −0.09 | 0.04 |
| *mdtH* | Efflux pump | 6 | 4 | 0.16 | −0.15 | 0.011 | 0.15 |
| *mdtL* | Efflux pump | 5 | 3 | −0.10 | −0.20 | −0.10 | −0.08 |
| *mefA* | Efflux pump | 13 | 9 | 0.08 | 0.02 | 0.02 | −0.07 |
| *oqxB* | Efflux pump | 13 | 17 | −0.02 | −0.04 | 0.06• | −0.12 |
| *pmrC* | Target alteration | 12 | 4 | −0.05 | −0.10 | −0.02 | 0.1 |
| *qacL* | Efflux pump | 16 | 18 | −0.12 | 0.32• | 0.11* | 0.22 |
| *silE* | Metal resistance | 15 | 4 | 0.17 | −0.04 | 0.19* | 0.07 |
| *tetA* | Efflux pump | 15 | 3 | 0.11 | −0.51 | −0.07 | −0.09 |

<sup>a</sup> Total number of samples = 16<sup>b</sup> Influent explanatory factors = Flow (m<sup>3</sup>/day) + Sewer pipe type (pressure main, gravity, both) + Design population size<sup>c</sup> Land explanatory factors = land cover + cattle + hospitals<sup>d</sup> Geographical explanatory factors = defined using PCNM eigenfunctions<sup>e</sup> Microbial community defined by eigen values obtained from PCoA using Bray-Curtis dissimilarity<sup>f</sup> Significance code- • p < 0.1, \* p < 0.05, \*\* p < 0.01, \*\*\* p < 0.001, values without notation were not statistically significant

A redundancy analysis was conducted to partition the variance in ARG sequence variant distributions among a number of explanatory variable groups (influent characteristics, land use, geographical location and microbial community composition; Table S4). A value of 0 indicates that the explanatory variables did not explain more of the variation than random deviations, whist a value of 1 explains 100 % of the variation. Negative values obtained indicate that the explanatory values explain less variance than a set of random normal deviates. Overall, the explanatory variables included in the redundancy analysis significantly explained only a small proportion of the variance in the distribution of ARG sequence variants. Influent factors (pipe design, influent flow and design population size) explained 30 % of the variance in the *mdt*G gene but did not significantly impact the diversity of other ARGs (Table 1). The geographical location explained only 11 % of the variation in the *qac*L efflux pump and 19 % of the variance in the *sil*E gene. Finally, the variation in ARG diversity explained by land factors (land cover, cattle numbers and hospitals) and microbial community composition was not statistically significant (Table 1). Taken together, these results suggest that sequence diversity is influenced by other parameters not included in the redundancy analysis.

### 3.3 Patterns of Sequence Diversity and Reported Hosts

For the purpose of demonstrating the value of this tool, four ARGs were chosen for further characterisation (Figure 2). The *bla*TEM and *bla*OXA genes, which confer resistance to beta-lactam antibiotics, were selected due to their clinical importance and detection in all sewer samples. The final two genes selected were *mdt*H (multidrug efflux pump) and *pmr*C (conferring resistance to peptide antibiotics) to study the impact of other mechanisms of resistance and to different antimicrobials. The ARG sequence diversity profiles between Gatineau and Quebec City (Figure 2a) for the remaining genes are reported in Figure S1.

**Figure 2.**
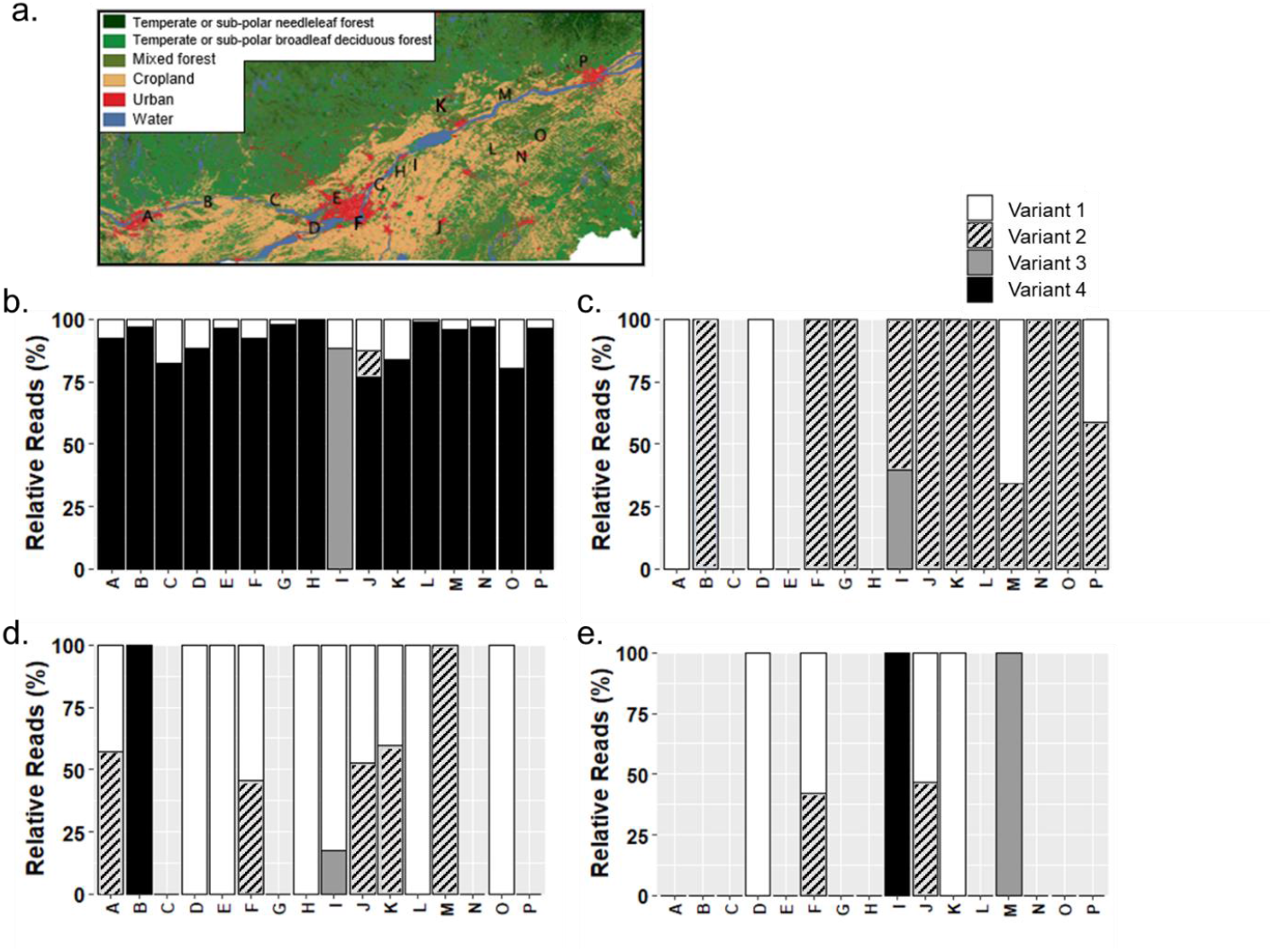
Sequence distribution for four genes detected in wastewater samples obtained in a transect along the Ottawa-St-Lawrence river basin between Gatineau (Point A) and Québec City (Point P). a) Sampling locations with respect to land coverage. Star indicates the island of Montréal. b) *bla*OXA c) *bla*TEM, d) *pmr*C, e) *mdt*H

**Figure 3.**
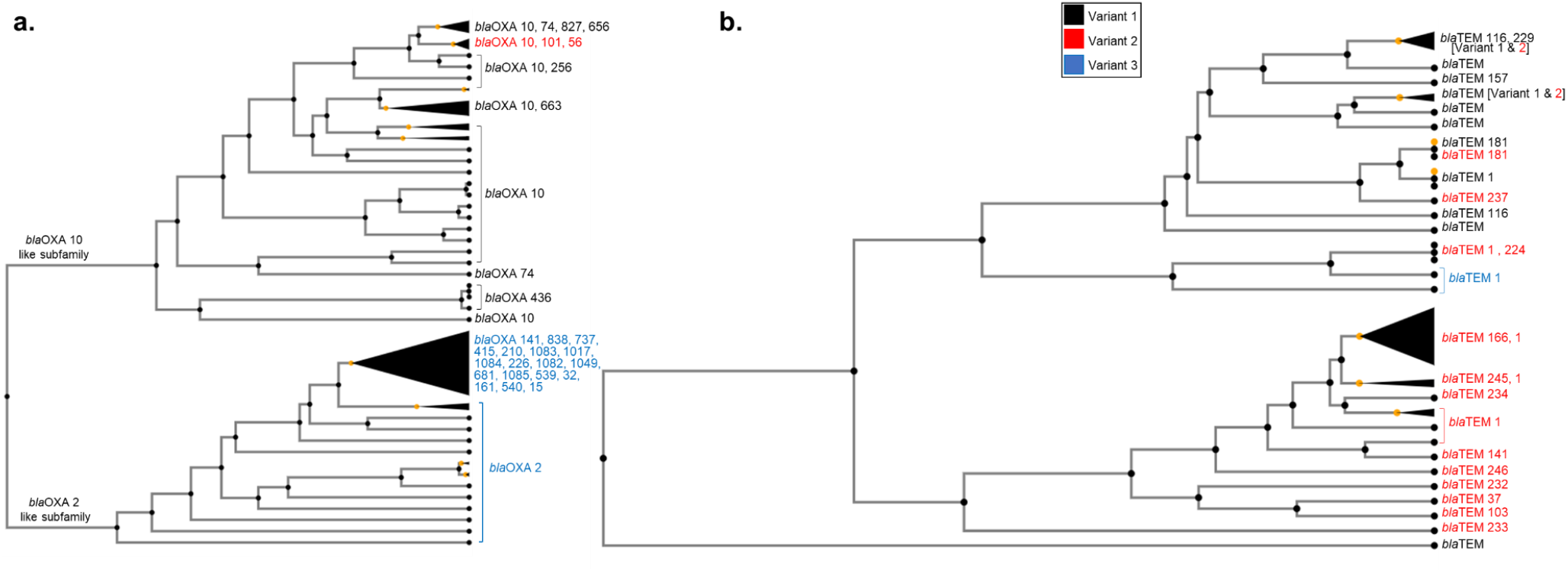
Multiple sequence alignment of ARGs containing each sequence variant in a) *bla*OXA and b) *bla*TEM. Sequences were aligned using MAFFT and the fast Fourier transform method (FFT-NS-2) (23) for multiple sequence alignment. Phylogenetic trees were generated using the average linkage (UPGMA) method.

Four variants of the *bla*OXA ARG were observed among the sewer samples (Figure 2b). The *bla*OXA Variants 1 and 4 were the most frequently detected, with a greater number of reads obtained for Variant 4 suggesting a higher abundance. The widespread detection of both Variant 1 and 4 suggest that the hosts are well adapted to the environmental conditions within the sewers or that there is a high influx of these variants from sources such as the human gut microbiome. Variant 2 and 3 were observed in only one sampling site each.

NCBI Blast (39) was used to gain more information about the previously reported hosts of each sequence variants. Based on the criteria that each variant may differ by only a single nucleotide polymorphism (SNP), only 100 % identity matches were accepted in this analysis. Variant 1, was previously detected in several hosts such as genus *Acinetobacter, Aeromonas* and *Enterobacter* (Table S6). Variant 2 was reported in fewer hosts. However, those identified largely overlapped with previously reported hosts of Variant 1 (Table S6). Surprisingly Variant 4 (most abundant and most frequently detected) corresponded to no hits with 100 % identity in the NCBI Blast database, whilst Variant 3 had 100 % identity over 100 % of its sequence with positive hits in over 30 species (Table S6). The closest reported sequence to Variant 4 matched 99 % of the query and was reported in hosts from the genus *Enterobacter, Klebsiella* and *Pseudomonas,* which overlapped with the genera reported to contain Variant 3. The lack of previous observation of *bla*OXA Variant 4 could be due to sampling bias of the database, which primarily includes clinical isolates and much fewer environmental strains. Conversely, Variant 4 differs from Variant 3 in only the last 6 nucleotides. Specifically, the last nucleotide of Variant 4 is different from that in Variant 3. Therefore, it is possible that this difference could be a sequencing/bioinformatics artefact, and Variant 3 and Variant 4 should be considered equivalent.

Among the sampled sewer communities, *bla*TEM Variant 2 was most prevalent (Figure 2c), whilst sub-dominant Variant 1 appeared to be present only in sampling locations typically far from Montreal. Both *bla*TEM Variant 1 and 2 have been reported in a large number of hosts (Table S6), suggesting that the gene is not restricted by phylogenetic barriers and possibly explaining the high occurrence of these variants among the sewer samples as some carriers are likely well adapted to this environment. *Bla*TEM Variant 3 has been reported only in synthetic constructs, which may justify its detection in only one wastewater samples and prevents any further hypothesis inference about its source. However, future sampling of different environments could enable tracking of such unique variants in different reservoirs to identify its origin and ecological vectors.

Four variants of the *pmr*C ARG were detected among the wastewater samples (Figure 2d). Variants 1 and 2 were the most frequently observed, and were reported in the same host species in the NCBI database (among genera *Escherichia, Salmonella* and *Shigella;* Table S6). Variant 3 and 4, which were identified in only one site each, have only been previously reported to be associated with *Escherichia coli*, suggesting a limited host range and consequently that they are less well adapted to persist in the sewer environment or the associated source microbiomes.

The *mdt*H gene was detected in fewer sewer samples, and the variants detected varied between sampling sites (Figure 2e). Variant 1 to 4 of the *mdt*H gene were all previously reported in taxa from the genera *Escherichia* and *Shigella.* The occurrence of all sequence variants in similar hosts somewhat explains the seemingly random assembly of variants in different locations, as it is likely that all are similarly adapted to the sewer environment.

Taken together, analysis of previously reported hosts using NCBI Blast suggest that the occurrence of a given variant may be influenced by the number of potential hosts and their ability to grow and compete within the sewer environment. Interestingly, many dominant variants returned no 100 % matches in the NCBI database. This may be due to the sampling bias associated with the database, where a greater number of samples are obtained from clinical than environmental settings. Given this presumed bias, this result raises questions on the interpretation of qPCR-based monitoring of AMR. These approaches provide little information on whether the quantified target is of any direct importance to the clinical outcomes in human or livestock, or if they merely represent general environmental circulation of genes in a given family. Such interrogations ask that we become increasingly precise in the identification of targets that are detected along with improving sample processing to expand the volume of environmental samples analysed.

### 3.4 Genetic context of ARG sequence variants

ARGs are common within the environment. However, their direct risk to public health is associated with the pathogenicity of their host and the likelihood of infection. This was aptly defined by Zhang et al., 2021, who outlined that the risk of a given ARG is largely dependent on 1) its enrichment in human associated environments 2) gene mobility and 3) its presence in pathogens. Mobile ARGs present in pathogens are considered to be current threats, whilst those that are mobile but not yet observed in pathogens are classified as future threats to public health.

To assess the mobility of each ARG variant, PLSDB (a plasmid database) was used to determine if they had been previously reported on a plasmid (Table 2) (40). The *bla*OXA Variants 1, 2 and 3 returned positive hits in the PLSDB, whilst no hits were obtained for Variants 4 (Table 2). Interestingly, plasmids containing *bla*OXA Variant 1 were detected in animals (dogs, pigs, chicken, duck), human and environmental (wastewater) samples, demonstrating that plasmids containing this variant have the ability to persist in a range of environments. Whereas Variant 2 and 3 were reported only on a plasmid obtained from a human sample.

**Table 2:** Occurrence of ARG sequence variants among 263 clinically relevant pathogenic bacteria reported in CARD

| Gene | Sequence Variant | PLSDB Database | Occurrence in Pathogens (n) |  |  |  |
| --- | --- | --- | --- | --- | --- | --- |
|  |  |  | NCBI Chromosome | NCBI Plasmid | NCBI WGS | NCBI GI |
| *blaOXA* | 1 | Y | Y (5) | Y (16) | Y (30) | Y (2) |
|  | 2 | Y | Y (1) | Y (1) | Y (2) | N |
|  | 3 | Y | Y (8) | Y (7) | Y (30) | N |
|  | 4 | NA | NA | NA | NA | NA |
| *blaTEM* | 1 | Y | Y (6) | Y (1) | Y (34) | Y (3) |
|  | 2 | Y | Y (28) | Y (35) | Y (54) | Y (11) |
|  | 3 | N | N | N | N | N |
| *pmrC* | 1 | N | Y (1) | N | Y (4) | N |
|  | 2 | N | N | N | N | N |
|  | 3 | N | N | N | N | N |
|  | 4 | N | N | N | N | N |
| *mdtH* | 1 | Y | Y (2) | Y (1) | Y (6) | N |
|  | 2 | N | N | N | N | N |
|  | 3 | N | N | N | N | N |
|  | 4 | N | N | N | N | N |

Variants 1 and 2 of the *bla*TEM gene also returned hits in PLSDB. However, Variant 3 was not previously reported on a plasmid. This suggests that short length sequences may be used to distinguish between mobile and non-mobile variants of a given ARG. Variant 2 was observed in plasmids associated with domestic animals (dogs), livestock (chicken, turkey, cattle and swine) and the environment (wastewater) again highlighting the potential for plasmids containing this variant to persist in different environments. Conversely, plasmids with 100 % match to Variant 1 were only reported in soil and feed additives, suggesting more limited movement. Based on the mobility of the variants, it could be hypothesised that *bla*TEM Variants 1 and 2 pose a greater risk to public health than Variant 3 due to their potential ability to transfer to pathogens and cause infections. Further studies on sequence variants associated with clinical infections and environmental strains could be used to expand our understanding of these patterns.

No 100 % hits in the PLSDB database were obtained for any of the *pmr*C variants, suggesting that this gene is likely less mobile than others and therefore poses less risk unless already present within pathogens. Of the *mdt*H variants, only Variant 1 returned positive hits in the PLSDB database. Variant 1 of the *mdt*H gene was the most frequently observed among the sewer samples, this could be related to the mobility of this gene and ability to transfer to hosts well adapted to survive within wastewater environments.

Considering the presence or absence of each variant in the plasmid database alone, it can be observed that the variable mobility of a given ARG could be detected using short read sequencing data obtained using multiplexed amplicon sequencing approaches. Mobile ARG observed in human associated environments have the potential to transfer into human pathogens and present a high risk of contributing to new multi-drug pathogens in the future. Mobile ARGs originating from clinical environments also have the potential to transfer to environmental bacteria, which could aid in the dissemination of AMR in the environment. This demonstrates the need for ARG sequence variant tracking, to monitor current and future high-risk sequence variants and develop effective control strategies to tackle the spread of AMR.

### 3.5 High Risk ARG Sequence Variants

The mobility of an ARG can be used as an indicator of the likelihood of future risk based on its ability to transfer into a pathogen. Whilst ARGs already present in pathogens present the greatest current risk due to the ability of the host to cause infection. The Comprehensive Antimicrobial Resistance Database (CARD) was used to determine the prevalence of the ARG sequence variants among the sequenced genomes, plasmids, whole-genome shotgun (WGS) assemblies and predicted genomic islands (GI) of 263 important pathogens. Results from this analysis were aggregated to provide the percentage occurrence of a given gene within each species (full method available (41)). For this analysis, the ‘perfect’ paradigm of RGI was selected which tracked perfect matches of each variant to reference sequences and mutations in the CARD database occurring within pathogens. This is consistent with the analysis conducted using NCBI Blast in Section 3.3 which considered only 100 % identity matches. This data was used to assess the prevalence and percentage occurrence of each variant among 263 clinically relevant pathogens to determine whether each ARG variant is a current risk to public health based upon available data.

Analysis of the *bla*OXA gene variants in CARD revealed that Variants 1, 2 and 3 have been previously reported in pathogens. Variant 1 and 3 have been previously reported on plasmids in several different pathogens, whilst Variant 2 was only identified on a plasmid in *Pseudomonas aeruginosa* (Table 2). Furthermore, Variant 1 has also been identified in the predicted genomic islands of two pathogens. In the three databases consulted thus far (NCBI Blast, PLSDB and CARD), no 100% identity matches have been obtained for both Variant 4, suggesting that multiplexed amplicon sequencing can be used to detect novel sequence variants. Given the detection of Variants 1 to 3 in pathogens, it could be suggested that these forms of the *bla*OXA gene present a current risk to public health due to the ability of the hosts to cause infection. The presence of Variant 1 and 3 in a greater number of pathogens also suggests increased likelihood of infection, and thus it could be hypothesised that they pose a greater risk to public health than Variant 2.

Both *bla*TEM Variants 1 and 2 were identified in pathogens using CARD. Variant 2 was identified in a greater number of pathogenic hosts than Variant 1, many of which were associated with plasmids demonstrating the mobility of *bla*TEM genes with this sequence variant. Variant 3 was not detected among the 263 pathogens included in CARD. Overall this alludes to *bla*TEM genes containing Variants 1 and 2 to being current threats to public health due to their previous associations with pathogens and their mobility. Conversely, Variant 3 has a lower chance of causing infection or transferring to pathogens because it was not previously reported in pathogens or on plasmids; thus, it should be considered a lower risk. These results highlight the different characteristics that can be inferred and monitored within a single ARG class and the need to study risk at sequence level.

To conclude this analysis,*pmr*C and *mdt*H were less frequently observed among the 263 pathogens in CARD. Variant 1 of the *pmr*C gene was detected in only four pathogens and was not associated with plasmids. Consequently, Variant 1 of the *pmr*C would be unlikely to transfer to other hosts via plasmid mediated horizontal gene transfer and thus is likely to remain associated with only a limited number of pathogens. Similarly, *mdt*H Variant 1 was observed in six pathogens. However only one report was on a plasmid. This suggests that the plasmid has recently transferred to pathogenic bacteria or limited mobility of the plasmid.

### 3.6 Wider Sequence Diversity of ARGs

Multiplexed amplicon sequencing was used to study the sequence variants within a targeted area of each ARG (approximately 275 base pairs). To gain more information about possible sequence diversity in the remainder of the gene and to inform future target design, NCBI Blast was used to obtain the full gene sequences in which the sequence variants have been previously reported. The obtained sequences were aligned to produce a phylogenetic tree enabling the diversity among the genes to be visualized.

Alignment of *bla*OXA sequences demonstrated that Variants 1 and 2 clustered together and belonged to the OXA-10 like subfamily of the gene, whilst Variant 3 and sequences similar to Variant 4 (as no 100 % identity matches were obtained) belonged to the OXA-2 like subfamily. Previous studies have shown *bla*OXA-10 to have weak carbapenemase activity, but variants of this gene within the subfamily such as *bla*OXA-656 are more efficient carbapenemases (42). ARG Variant 1 was identified in numerous variants of the *bla*OXA-10 gene, many of which such as *bla*OXA-17 and *bla*OXA-74 display extended spectrum beta lactamase activity (43). Whilst *bla*OXA sequence Variant 2 was identified in only a small cluster of the *bla*OXA-10 subfamily, that included the *bla*OXA-101 and *bla*OXA-56 genes which display narrow spectrum beta lactamase activity (44). The *bla*OXA-2 like subfamily which clustered separately (containing Variants 3 and 4 of the current study), have been demonstrated to have more broad-spectrum activity. In the future, primer design could be optimized to target specific subgroups within the *bla*OXA-2 family and enable a more accurate assessment of ARG activity and hence risk.

Multiple sequence alignment of the *bla*TEM ARG showed the sequence variants to be more widely distributed among different classes of *bla*TEM genes. The amplified sequence variants were often identified in the same *bla*TEM genes. For example, Variant 1 to 3 were present in *bla*TEM-1 and Variant 1 and 2 are both in *bla*TEM-181. Furthermore, the variants could not be used to distinguish between genes with narrow or extended spectrum activity. Among the *bla*TEM genes, extended spectrum beta-lactamases such as *bla*TEM-181 and *bla*TEM-166 are derived from genes with narrow spectrum beta-lactamase activity such as *bla*TEM-1 by mutations which alter the enzymes active site (45). In the future, primer optimization to target these mutations could improve risk assessment by providing additional information on differences in phenotypic resistance.

### 3.7 High Risk ARG Sequence Variants

The risk and genetic context analyses conducted thus far have relied upon data available in public databases, which are often bias towards clinically derived isolates. To further investigate the genetic context of the specific sequence variants in the samples collected in the current study, an inverse PCR approach was used to obtain sequence information on the flanking regions of the ARG. For this analysis, we focused on the *bla*OXA-2 like variants (variant 3 and 4) as they were the most widely detected among the sewer samples. Clones obtained from 3 samples from along the transect of sampling sites (site A, F and P) were analyzed. Using this approach, it was found that all of the flanking regions sequenced were previously reported to be associated with mobile genetic elements including integrons such as *Int*I1, transposons such as tn21, and plasmids. This suggests that mobility may be crucial to the persistence of ARG sequence variants in the environment. Among the plasmids, some such as pALT31 and pALT33 were present in all three sampling sites. These plasmids were first reported in wastewater biosolids, and have been demonstrated to harbor ARGs to numerous classes of antimicrobials and also persist in the absence of antibiotics (46). In the future, inverse PCR approaches could be used in combination with multiplexed amplicon sequencing approaches to gain further information on the hosts of ARGs in the environment and confirm the findings of database focused analyses.

### 3.8 The Future of Multiplexed Amplicon Sequencing

Using targeted amplicon sequencing, this study identified novel ARG sequence variants not previously reported in NCBI, CARD or PLSDB databases. Information on ARG sequence diversity can be used to infer the risk of a given variant based on its mobility and occurrence within pathogenic bacteria. As previously discussed by Zhang et al. 2021, we demonstrate that different variants within an ARG class present different risks. This information could be a valuable asset in environmental surveillance and could be used to inform management strategies to control emergence and spread of AMR.

Targeted amplicon sequencing is a relatively low-cost technique, which allows multiple samples to be analysed in a single sequencing run with a detection limit much lower than achievable by other methods. This will allow a greater number of samples to be analysed and patterns in sequence diversity to be observed. In the future, targeted sampling of AMR hotspots could be used to build databases of unique and common ARG sequence variants observed in each environment. This data would be beneficial in environmental studies for AMR source tracking, and in clinical settings to determine the risk of a given variant or the mechanistic origin or recurrent nosocomial infections.

