## Supplementary Material for "Multiplexed Amplicon Sequencing Reveals High Sequence Diversity of Antibiotic Resistance Genes in Québec Sewers"

| Table S1: Sample ID of Sewer Samples and Sequences Obtained | | |
| --- | --- | --- |
| Sampling Location | Given ID | Sequences Obtained |
| Gatineau | A | 92066 |
| Granby | J | 86674 |
| La Prairie | F | 69821 |
| Lachute | C | 82091 |
| Papineauville | B | 75985 |
| Pincourt | D | 104849 |
| Plessisville | O | 97617 |
| Rosemere | E | 101762 |
| Shawinigan | K | 92524 |
| St Casimir | M | 96397 |
| St Eulalie | L | 65885 |
| St Ours | H | 87912 |
| St Robert | I | 87473 |
| Stoneham | P | 92964 |
| Verchères | G | 63155 |
| Victoriaville | N | 82945 |

| Table S2: Colour scheme applied for mapping land use | | |
| --- | --- | --- |
| Value | Description | Colour Scheme |
| 1 | Temperate or sub-polar needleleaf forest | RGB 0 61 0 |
| 2 | Sub-polar taiga needleleaf forest | RGB 148 156 112 |
| 3 | Tropical or sub-tropical broadleaf evergreen forest | RGB 0 99 0 |
| 4 | Tropical or sub-tropical broadleaf deciduous forest | RGB 30 171 5 |
| 5 | Temperate or sub-polar broadleaf deciduous forest | RGB 20 140 61 |
| 6 | Mixed forest | RGB 92 117 43 |
| 7 | Tropical or sub-tropical shrubland | RGB 179 158 43 |
| 8 | Temperate or sub-polar shrubland | RGB 179 138 51 |
| 9 | Tropical or sub-tropical grassland | RGB 232 220 94 |
| 10 | Temperate or sub-polar grassland | RGB 225 207 138 |
| 11 | Sub-polar or polar shrubland-lichen-moss | RGB 156 117 84 |
| 12 | Sub-polar or polar grassland-lichen-moss | RGB 186 212 143 |
| 13 | Sub-polar or polar barren-lichen-moss | RGB 64 138 112 |
| 14 | Wetland | RGB 107 163 138 |
| 15 | Cropland | RGB 230 174 102 |
| 16 | Barren lands | RGB 168 171 174 |
| 17 | Urban | RGB 220 33 38 |
| 18 | Water | RGB 76 112 163 |
| 19 | Snow and Ice | RGB 255 250 255 |

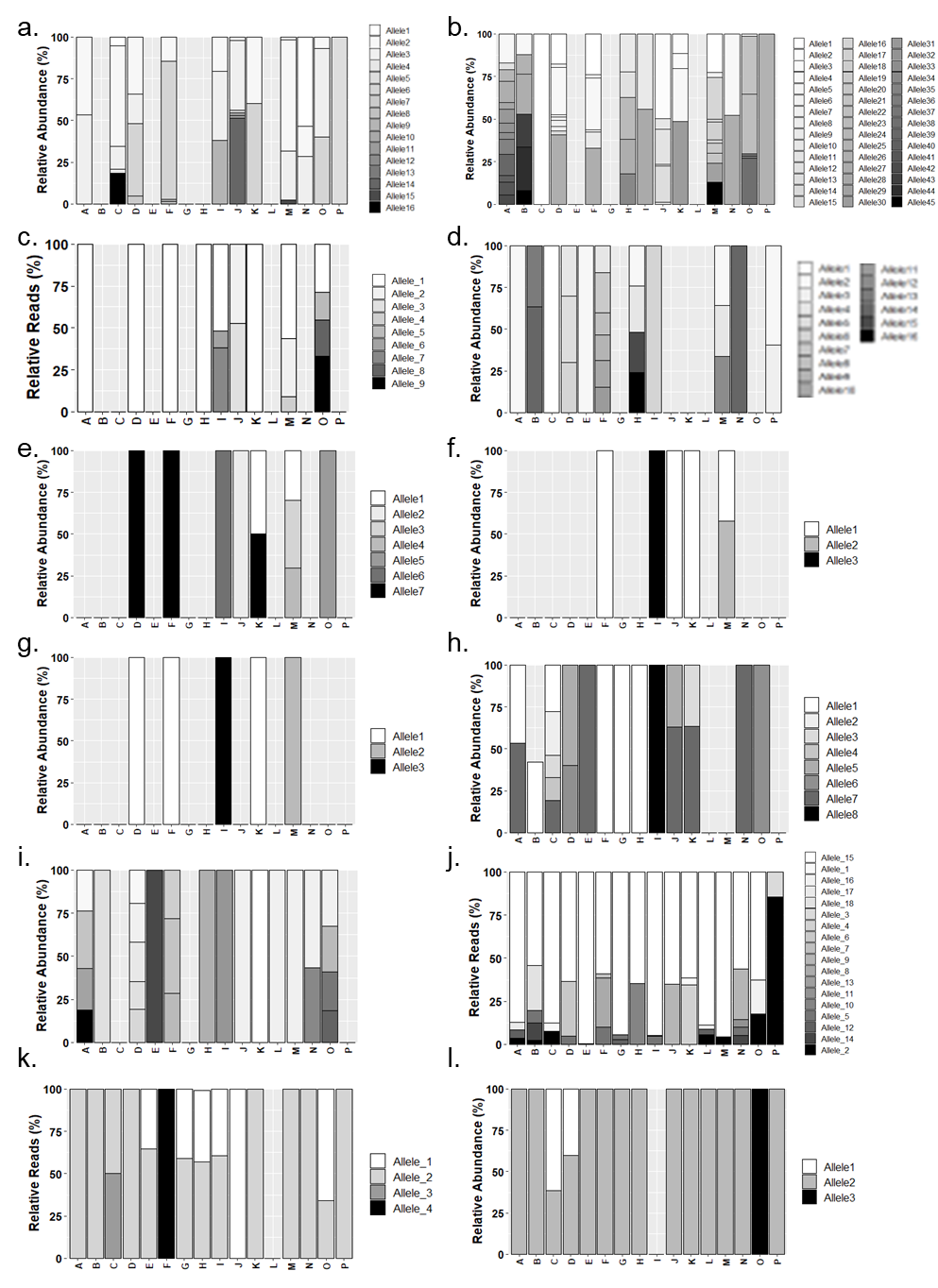

Figure S1- ARG sequence diversity of a) arsA b)blaACT c) blaSHV d) copA e) mdtF f) mdtG g) mdtL h) mefA i) oqxB j) qacL k) silE l) tetA

| Table S3 Multiplexed Amplicon Sequencing Primers | | | |
| --- | --- | --- | --- |
| Forward Primer Name | Forward Sequence | Reverse Primer Name | Reverse Primer Sequence |
| ampC | CGCCGTAATTCAGCCGCTGA | ampC | GGAGGTGTGGGTCGCCAAATTC |
| ampCblaDHA | TAAACCGCTGATGGCACAGCA | ampCblaDHA | GCACCCAGCACACCTGTGA |
| amra | GCAGCAGGACGTCGAGGTG | amra | TTTCATCTGCGTGGCCTCGAC |
| ant2D | CGTGTAACCCGCAAGCACGATGATA | ant2D | AGCCTGTAGGACTCTATGTGCTTTGTAGG |
| ant3D | TGGCAGCGTAAGGACATTCTTGC | ant3D | ATTGCCCAGTTGGCAACGA |
| ant6ia | GTTCGCCATGAGCTGCTGA | ant6ia | ATATCAGCGGCATATGTGCTATCC |
| ant9 | AGGATATTGTCCCTTGGCATTTCC | ant9 | AGATGCTGTCAGCCACATTCTG |
| aph2Id02aph901 | ATTCATGCGCCCAATGGTTTCA | aph2Id02 | TGCCCGCATGTATATCAGAATGAC |
| aph3ia | TGAAACATGGCAAAGGTAGCGTTGC | aph3ia | TCGTGATTGCGCCTGAGCGA |
| aph3via01 | GCTTGCTATCTATAAGGAGGCACTCA | aph3via02 | CACGAGTCTCGGTTAACTCATTCC |
| aph4ia | GCGTGGATATGTCCTGCGGGTAA | aph4ia | GACCGATTCCTTGCGGTCCGA |
| aph4ib | GTTCGTCCACGGCGACCTG | aph4ib | CAGAGGAACTGCGCCAGTTCC |
| aph6ic01 | ACCAGGCGCTGCTGATGG | aph6ic01 | GCTCGCCGAAGTCGAGCA |
| aph6ic02 | GCTGCTCACCACCCACTCG | aph6ic02 | CTGGAACCATTCCTGTAGCGGATG |
| ARR01 | TTTGGCGATTGGTGACTTGCTAACCAC | ARR01 | CCACGGCGCTTTAAGTCCTCCAAC |
| arsA01arsA03a | CGATGGCCGGGCTGGAAA | arsA01 | AGTAATAGGGTATCTGTCGGTAGCTC |
|  |  | arsA02 | GTAGTAATAGGGTATCTGTCGGCAACTC |
| arsA02arsA03b | CCAATGGCCGGGCTGGAAA | arsA03aarsA03b | GGAGTAACAGGGTATCTGTCGGTAGCTC |
| betompK | ACCGGTTACGGCCAGTGGGA | betompK | GCAAAGTTCAGGCCGTCAACCAGA |
| bl1ec | GGTATGGCGGTAGCGGGAATTTATCA | bl1ec | TCGCTTGAGGATTTCACCTCATCCG |
| bl2akcc | CGTACTGCGCGACCTCGAC | bl2akcc | GCTCGCCGGAGTTCAGCTC |
| bl2anps | CGAGCCGTCTCGACCGCATC | bl2anps | CCGAGATGAACATGGTCGCGATCC |
| bl2bfona02 | CAGCGCGATTCCCGGCGATA | bl2bfona02 | CATCCTGCTGCGGCTGGGTAAA |
| bl2btle01 | ATGTCGTTGGCCGAGTTGTG | bl2btle01 | GGTCTGGTTGGCCCGCATCC |
| bl2btle02 | CAACACCGCCGGCAACCTG | bl2btle02 | CGGTCTTGTCGGCCGTGGTC |

| Table S3 Contunued | | | |
| --- | --- | --- | --- |
| Forward Primer Name | Forward Sequence | Reverse Primer Name | Reverse Primer Sequence |
| bl2bula01 | CCGCCATCCCTGGCGATGA | bl2bula01 | TGCTGTGGCTGTGTAAAGTAGG |
| bl2bula02 | TGTGCAGCACCAGCAAGGTGATG | bl2bula03 | GCCGCCAAGATGCTCAAGGA |
| bl2bula03 | TCAGCAGCAACTGTCTGAACTGG |  |  |
| bl2cbro02 | TGTGTTGGCCTCATCTGCACATC | bl2cbro02 | GCCAGGTGGCACCCTTTCC |
| bl2dicrNPS01 | ATCCTCGTGCGCAACAACGATAC | bl2dicrNPS01 | GGAGAAAGGCAACCTGTTCTTCCG |
| bl2dicrNPS02 | AAGAGCGCATTCAGGGTCAGCTG | bl2dicrNPS02 | AGCCGCCCAACCGGTCTTG |
| blaACI | CCTGCGCGAAATGAACCTGA | blaACI | CGTGCGGCGCGACATTA |
| blaACT01 | TGGCAGCCGCAGTGGAA | blaACT01 | GGCGTAATGCGCCTCTTC |
| blaACT02 | CCGGATGAGGTCACGGATAACGC | blaACT02 | TCCCAGCCCAGACCCTGATACA |
| blaACT03 | TGGCGCAGTCTCGCTACTGG | blaACT03 | GCCACGTAGCTGCCAAACC |
| blaAMPH | TGATGCGCTGGATGCAGCAG | blaAMPH | CCAGGTCGTTGATGCCATCGC |
| blaBEL | ATGGCACAGACTGTGTCAAAGCTG | blaBEL | GACGTGCTAACTTTGCTACAGAAGCA |
| blabjp1 | CGAGCTGGGAGATGACCGTCAAG | blabjp1 | GCTTGGCGAGCTGCTTGTCG |
| blablaec | CGCCGAACAAACCGATACTGGA | blablaec | CGTCACACAGCAAACGTTGTTTGA |
| BlaCAU | CTGCACCAGCTGGACCAC | BlaCAU | TGTTGAAGGCGGCTTCGG |
| blaCMY01 | CTAACTCCAGCATTGGTCTGTTTGG | blaCMY01 | GGCCAGTTCAGCATCTCCCA |
| blaCMY02 | ACTGGCAGCCGCAGTGGA |  |  |
| blacphA | TGCTGGAGGTGATCAACAACAACTAC | blacphA | CTCCTTGAGGATGCAGTTGCCA |
| Blactxm01 | GGAAGTGTGCCGCTGTATGC | Blactxm01 | AAGCGCTCATCAGCACGATAAAG |
| Blactxm02 | AAAGCTGGCGGCGCTGGA | Blactxm02 | ATTGTCGCTGTACTGCAACGC |
| Blactxm03 | CCTTCCGTCTGGACAGAACCG | Blactxm03 | GAATGCTCGCGCTACCGGTA |
| blaFOX | GCCGATGATCAAGGAGTATCGG | blaFOX | GGGAACTGCAGCGGCAA |
| Blaimp01 | CAGCACGGGCGGAATAGAGTG | Blaimp01 | GCGGACTTTGGCCAAGCTTCTA |
| blaLAT | GAGTTACGAAGAGGCAATGACCA | blaLAT | ACATATCGCCAATACGCCAGTAG |
| blaLCR | GTGCAGAGAACCAGGCAATCGC | blaLCR | TGTCTAGAGTCTGGTCCTGGTTCCAA |

| Table S3 Continued | | | |
| --- | --- | --- | --- |
| Forward Primer Name | Forward Sequence | Reverse Primer Name | Reverse Primer Sequence |
| blaMIR | AGGCCATTCCGGGTATGGC | blaMIR | GGTTGCCAGATCCAGCATGC |
| blaMOX | ACCAGCTCGGCGGATCTG | blaMOX | GAGCCGGTCTTGTTGAAGAGC |
| blaOXA03 | AGGCACGATAGTTGTGGCAGAC | blaOXA03 | GTAGAATTCCGCATTGCTGATCGC |
| blaOXA04 | GACGGAAAGCCAAGAGCCATGAA | blaOXA04 | CACCATGCGACACCAGGATTTGA |
| blaPAOPDC | TACAACCGGTGATGAAGGCCTATG | blaPAOPDC | GCGGTATAGGTCGCGAGGTC |
| blaPER | TCGCAAATGAAGCGCAGATGCA | blaPER | CTCTGGTCCTGTGGTGGTTTCGA |
| blaPSE | CGGACCTGCTGCAATTCATTGCC | blaPSE | ACACTACATAGGCGCCGAAACCG |
| blaSHV11LEN01 | GCCGCCATTACCATGAGCGA | blaSHV11LEN01 | GCACGGAGCGGATCAACGG |
| blaSHV11LEN02 | GCCGCCATTACCGTGAGCGA | blaSHV11LEN02 | AGCACGGAACGGATCAACGG |
| blaSHV11LEN03 | CGCCGTCATTACCATGAGCGA | blaSHV11LEN03 | GCACGCAGCGGATCAACGG |
| blaTEM | GAACCGGAGCTGAATGAAGCC | blaTEM | CGGGAGGGCTTACCATCTGG |
| blaVEB | CCATTTCCCGATGCAAAGCGTTA | blaVEB | GGGAATTCCTCTTTAATCGGACTCCA |
| blaVIM | TCTCCACGCACTTTCATGACGAC | blaVIM | GTCGGTCGAATGCGCAGCAC |
| catA1 | CGACGATTTCCGGCAGTTTCTACACATA | catA1 | CCCTGCCACTCATCGCAGTACTG |
| chlchiA | AGCTCTTGTTTGGACCGCTATCG | chlchiA | CATGCCCAAACGTAGAAACGCAAA |
| cmlA101 | CAACGAGTTGCGGGCTTGCA | cmlA101 | TGGCAAGCAATACTGCTCCAGCTA |
| cmxA | GCGTGCACTGTCGATCCTGCTC | cmxA | GCCAGGAAGGTGAATGCCGCAA |
| copA | GCGAGGTCGGTTGGGCTCA | copA | AGAACGGCACCTACTGGTACCAC |
| dfhr01 | GTGCCAAAGGTGAACAGCTCCTG | dfhr01 | CCACCACCTGAAACAATGACATGATCC |
| dfhr02dfrA1,5,14 | GCGGTCCAGACATATCCTGGTC | dfhr02dhfr02 | CCAGACACTATAACGTGACCGTTGA |
| dfrA1201 | GGTAATCTCAAGCCAAGCTAACTACC | dfrA12-01,04,04a | ACGCGCATAAACGGAGTGG |
| dfrA1202 | AATCTCACGCCAAGCTAACTACC | dfrA12-02,03,204b | AATGGAGTGGGTGTACGGAATTACAG |
| dfrA1203adfrA1203b | GGTAATCTCACACCAAGCTAACTACC |  |  |
| dfrA1204adfrA1204b | TGGTAATCTCACGCCAAGCTAACTACC |  |  |
| dfrA22 | GTCGTTATGGGCCGCAAGACG | dfrA22 | GTGCGTGTACGTAATTGTGGCTTGA |
| Table S3 Continued | | | |
| Forward Primer Name | Forward Sequence | Reverse Primer Name | Reverse Primer Sequence |
| dfrA2501 | GTGATCGGTTGCGGTCCACAC | dfrA2501 | AGCGTAGAGGCCATGGGCAA |
| dfrAq | ACATACCCTGGTCCGCGAAAG | dfrAq | CGCCACCAGACACTATAACGTGA |
| dhfr01 | CGGGAATGGCCCTGATATTCCATGGA | dhfr01 | CACCACCTGAAACGATGACATGATCCG |
| emrD | GATCCCGGCGGCGTTCTTC | emrD | CGGCAGCATCGCCGAAAGC |
| ereA01 | GAAACGGCTCGGCAGGAG | ereA02 | ATTGTTGTGCGCCAGCAGA |
| ereA02 | GGAAACGGCTCAGCAGGAG | ereA01 | TGATTATTGTGCGCCAGCAGA |
| ereB02 | CAGCTCATCGATCACCTCATGAAACCG | ereB02 | CACGTACGGAAGTATCTCCCTCAA |
| lncWtrwB | TTGTGGCGGGCCTGCAATCG | lncWtrwB | AGGCGGTGAGGTCGGGCAA |
| macA | GACGCGCTGCGTAATGTTGA | macA | TTGCGTCTGTTTCACAGCCATC |
| marA | CCAGACGCAATACAGACGCTATTACC | marA | TTCGCCTTGCATATTGGTCATCCG |
| marR | CACAGTTTAAGGTGCTCTGCTCTATCC | marR | GCAAATACTCAAGTGTTGCCACTTCG |
| mcr101mcr102 | GCTATTAATCATGGGCGCGGTGA | mcr101 | GGCATGATCGGATTGACATACGAACGC |
| mdtF | TTTGGTTCCGAGTACGCCATGC | mdtF | GTTATAGCGCGCCACGGTGGA |
| mdtg | AAGAGATGCTGCATATGCGGGAAG | mdtg | TGGCGTCTGAACGTATGACATTGG |
| mdth | CTGCCGTTAAATGGATGTATGCCA | mdth | CCAATCGCCAGAACCAGACG |
| mdtL | CCAGCGAGGCGCAGTTGCATA | mdtL | GCTAACACCGGAATGATGCAGGTA |
| mefA | GATCTGCGATAGTCTTGTCTATGGC | mefA | ACTGACTATAGCCTGCACATTTCG |
| mexD | CTGGTGATGTTCCTGTTCCTACAG | mexD | GAAGGCCAGCGGCAGGAACAC |
| mexE | GACACGTCGTCCGGCACGA | mexE | GCTGCGTGCCGTTCACGAC |
| mgt | TCGGTTCGGCCTTCACCAA | mgt | GGCGTTGCCGAACTGGTC |
| mphA01 | GCTGGCTCGACGACGATTC | mphA02 | CCGAGTCGAGGGCGAAGA |
| mphE | AAGTGAGCAATTGGAAACCCGCTA | mphE | AGGCCGCTGCTCTTTCTAAAGTC |
| msrD01 | GGCAAGCTAGGTGTTGAGCAATTAG | msrD01 | CCTTCACGATCTAAATGGCTCGTA |
| msrD02 | GCCTTATCGGCACAGGTTCATGG | msrD02 | TTCCTTCACGGTCTAGATGACTGGTA |
| msrD03 | GGCAAGCTAGGTGTTGAGCA | msrD03 | TCCTTCACGGTCTAAATGGCTCGTA |
|  |  | nimA01 | TCCTGTCATGTGGTCGATGCA |

| Table S3 Continued | | | |
| --- | --- | --- | --- |
| Forward Primer Name | Forward Sequence | Reverse Primer Name | Reverse Primer Sequence |
| nimA02 | ACAAGCTGGACGCCATCGC | nimA02 | AGCTCGATGGCCTCTTTGCC |
| obrj | CGCGCTGGACTACCAGGAAGC | obrj | AGGCTGATGCGCGGGAAGAAC |
| okp | AACACCTTGCCGACGGGATG | okp | ACCATCCACTGCAGCAGCTG |
| oprn | CGCTTCGAAAGCCTGTGGTGGAA | oprn | AGGGCGTCGCTGGACTCCA |
| oqxB | TGCTGGTGGTGCTGGTAGTGATC | oqxB | AAACGCCATCGGCACGAACAC |
| otra | GTACGGGTGCGCTTCGAC | otra | GACTGCGAACGCCGGTAC |
| PBRT | GCAGGAAGGCCGTCAGATCAC | PBRT | CTCATGGCGCTCTATCAAGTCGTC |
| pcoA | GCCATTCCGGTCTGCAGGA | pcoA | GGCCTGCCCGTTCATGAGA |
| pilDPA | CGACCTACAACCTGGTGCTG | pilDPA | GGTGATCGGCATCGATCAGG |
| PmrC01 | ACGCCATCACCTTGCAGGTTA | PmrC01 | GGCAGTTTCAGTACCGTGCATG |
| PmrC02 | TTCGGCAGACAATAACCACCATAACC | PmrC02 | TATTGTCATCGGTGAGCGTTCAGAAC |
| qacFH01 | GTTGCAATCTTTGGCGAGGTCA | qacFH01qacFH03a | TTTAGAACGGCGACACCACTG |
| qacFH02 | GCTGTTTCAATCCTTGGCGAGGTCA | qacFH02qacFH03b | CGCTGACCTTGGATAGCAGGTTTAGAAC |
| qacFH03aqacFH03b | CTGTTTCAATCTTTGGCGAGGTCA |  |  |
| qnrB01 | CGACCTGAGCGGCACTGAATTTA | qnrB01 | GCTCGCCAGTCGAAAGTCGAA |
| QnrS1S3S5 | CTGCAAGTTCATTGAACAGGGTGA | QnrS1S3S5 | CACCTCGACTTAAGTCTGACTCTTTCAG |
| qnrS2 | ATGCCAGCTTGCGATGGCAAA | qnrS2 | GTGGCATAAATTAGCACCCTGTAGGC |
| QnrVC4VC5VC701 | TGGTATCGAGTTCAGAGAATGCGA | QnrVC4VC5VC702 | AACCATACAACTCCGAGTGGCTCAA |
| robA01 | TCAAATGCGCGTGCAGTTCTGG | robA01 | GTAGCGCTCAATATCCTGACCTTTAC |
| satG | CCGGGACTTACCGTGGGTTC | satG | CTTCCAGGCATCGGCATCTCA |
| silE | TCATGGGCAACCGCTGCTCTTTC | silE | GGGCCACTGAAACCGTGAATATCCATG |
| spcN | TGCTGCTGCGCGAGCA | spcN | CACGTCCAGCAGGCTCCAG |
| strA | CGGCAATTCCGGGAGTACCG | strA | CCAGTTCTCTTCGGCGTTAGCAA |
| sul1 | GACGAGATTGTGCGGTTCTTCG | sul1 | ATTTCGCGAGGGTTTCCGAGA |

| Table S3 Continued | | | |
| --- | --- | --- | --- |
| Forward Primer Name | Forward Sequence | Reverse Primer Name | Reverse Primer Sequence |
| sul201 | TATCGCGCCGGTGCTGGA | sul201sul202 | GGACAAGGCGGTTGCGTTTG |
| sul202 | GTATCGCGCCGGTACTGGA |  |  |
| sulIII | TGATACAACTGAAGTGGGCGTTGTGGA | sulIII | CACGCTTTACACCAGCCTCAACTAAAGC |
| surA1AB | GCTGGCGATCATAACTATTCTGACTC | surA1AB | CGCTTGTTTAGCGTGTTCCCA |
| tcmA | CCATCGTGGCCATAGCCAAC | tcmA | CGATCGCCATGTTGAGCTTCTCG |
| tcr3 | ATCGGCGTCGGCATCTTCGG | tcr3 | GTCTGGGACAGGCCGATGCC |
| tet39 | TGACTGAGAAGGTTCAGGAGCAATC | tet39 | ACCAATTGCATCGCAAGCTATTCC |
| tet40 | CTGCTGGGAAACATAAACCTGCAA | tet40 | CCGAACCAAGTGACGCTGTTC |
| tetA | TCGGCGAGGATCGCTTTCACTG | tetA | TCCTCATCCACCTGCCTGGACA |
| tetB01 | GCCAGTCTTGCCAACGTTATTACG | tetB01 | ATCCCTGAAAGCAAACGGCCTAA |
| tetE | TGATTGCTGGACCAGTCATTGG | tetE | CCATACGAAGCGCTCTTCTCC |
| tetG | TTCAAGCCGGCTTGGAGAGC | tetG | ACAATCCAAACCCAACCGTTCCA |
| tetL01 | CTGGGTGAACACAGCCTTTATGTTAACC | tetL01 | GGCTATCATTCCAACAATCGCTGGAC |
| tetO | GCAGGGACAGAACTATTAGAGCCATATC | tetO | GCTAACTTGTGGAACATATGCCGAAC |
| tetQ | TGGATTGAAGACCCGTCTTTGTCC | tetQ | AGCAGGTGTACTTACCGGGCTATA |
| tetR | TGCTCGACGCCTTAGCCATTG} | tetR | GCGACTTGATGCTCTTGATCTTCCAA |
| tsnR | GCGGTGCAGCGGATCATCGA | tsnR | CCGTCGAGGACGACGACGTC |
| vanA | GTTAAGCCGGCGCGTTCAG | vanA | TTTGGTCCACCTCGCCAACA |
| vanSB01vanSB02 | CATAGGGCGCACGGTTGC | vanSB02 | TCGCGCCTGATCAATGCG |
| vanXA01 | GTGGACGGCTACCTGGTGAACC | vanXA01 | CCCATGTCGGCGAGCTCACC |
| vph01 | GAGTTGTTCCCGCTCATGTCC | vph02 | CCGAGGTCGCCATGGACCA |
| vph02 | GCCGTACCTGGTGCTGAGC |  |  |
| mfpa | CGTGAATCTGGCCGAGTCACAAC | mfpa | CGCAAGTCGGCGTCATCCAG |
| norA | ACCAGGGATTGGTGGATTTATGGC | norA | CGCCACCCGTAATAGCAATCGA |
| qacAB01 | GGAATTGGAATTGCAGCCATTGGC | qacAB01qacAB02 | CGCCCACTACAGATTCTTCAGCTAC |

| Table S3 Continued | | | |
| --- | --- | --- | --- |
| Forward Primer Name | Forward Sequence | Reverse Primer Name | Reverse Primer Sequence |
| qacAB02 | AATTGGAACTGCAGCCATTGGC |  |  |
| qnrB01 | CGACCTGAGCGGCACTGAATTTA | qnrB02 | TCCCACAGCTCGCATTTCTCCA |
| qnrB02 | AAACGGCAGCTTTATGCTGTGTGA | qnrB01 | GCTCGCCAGTCGAAAGTCGAA |
| qnrD | GACAGGAATAGCTTGGAAGGGTGTG | qnrD | ACGGCGCCAGTTATCACAGTG |
| qnrVC01 | GCTCAAACCTTCGAGATACACAGTTC | qnrVC01 | CCTCGAAGATTTGCACCAATCCATC |
| qnrVC02 | TGGTATCGAGTTCAGAGAGTGCGA | qnrVC02 | CATACAACTCCGAGTGGCTCAAATCA |
| QnrVC1VC3VC6 | GGGCACTAGAAGGGTGCGA | QnrVC1VC3VC6 | CCTCGAAGATTTGCACCAATCCA |
| QnrVC4VC5VC701 | TACAGCTCCGAGTGGCTCAA | QnrVC4VC5VC701 | TGGTATCGAGTTCAGAGAATGCGA |
| QnrVC4VC5VC702 | TTGGTATCGAGTTCAGAGAGTGCGA | QnrVC4VC5VC702 | AACCATACAACTCCGAGTGGCTCAA |

| Table S4: Sample metadata used in partitioning of variance | | | | | | | | |
| --- | --- | --- | --- | --- | --- | --- | --- | --- |
| ID | Sample | Longitude | Latitude | Design Pop. | Avg. Flow (m³/day) | Cattle/Division^a^ | Land Cover^b^ | Sewer Pipe Type |
| A | Gatineau | -75.6047 | 45.4744 | 230000 | 143708.6 | High | Urban | Gravity |
| B | Papineauville | -75.0217 | 45.6122 | 1700 | 1724.8 | Medium | Forest/Cropland | Force |
| C | Lachute | -74.3672 | 45.6306 | 12809 | 10998.8 | Medium | Forest/Cropland | Combined |
| D | Pincourt | -73.9872 | 45.3572 | 13545 | 7660.9 | Low | Semi-Urban/Cropland | Force |
| E | Rosemère | -73.7694 | 45.6497 | 27000 | 22186.1 | Low | Urban | Force |
| F | La Prairie | -73.5553 | 45.4153 | 64430 | 61073.3 | Low | Urban/Cropland | Gravity |
| G | Verchères | -73.3536 | 45.7803 | 3600 | 4343.0 | Low | Semi-Urban/Cropland | Force |
| H | St-Ours | -73.1511 | 45.9042 | 2055 | 1270.2 | Low | Forest/Cropland | Force |
| I | St-Robert | -73.0066 | 45.9765 | 376 | 113.8 | Low | Semi-Urban/Cropland | Gravity |
| J | Granby | -72.7736 | 45.3706 | 38400 | 56084.1 | Medium | Forest/Semi-Urban/Cropland | Gravity |
| K | Shawinigan | -72.7561 | 46.5539 | 24925 | 19506.8 | Low | Forest/Semi-Urban/Cropland | Force |
| L | St-Eulalie | -72.2556 | 46.1281 | 450 | 392.7 | Low | Forest/Cropland | Force |
| M | St-Casimir | -72.1356 | 46.6572 | 1265 | 808.8 | Low | Forest/Cropland | Force |
| N | Victoriaville | -71.9764 | 46.0511 | 34125 | 33942.6 | Medium | Semi-Urban/Cropland | Force |
| O | Plessisville | -71.7897 | 46.2578 | 8000 | 5609.2 | Medium | Forest/Semi-Urban/Cropland | Force |
| P | Stoneham | -71.3756 | 46.9775 | 3780 | 1288.4 | Medium | Urban/Forest | Force |
| a: Number of cattle in division where low= 0-2001, medium=2001-5871 and high= 5871-543566 | | | | | | | |  |
| b: Dominant land use in the division | | | | | | | | |

| Table S5: Significance of environmental variables obtained using function envfit permutation test in R Vegan Package | | | |
| --- | --- | --- | --- |
| Variable | r^2^ | Pr (>r) | Significance^1^ |
| Longitude | 0.0573 | 0.682 | NS |
| Latitude | 0.1175 | 0.449 | NS |
| Population Design | 0.0453 | 0.690 | NS |
| Volume/year m3 | 0.0874 | 0.489 | NS |
| Sewer Type (Combined) | 0.1637 | 0.287 | NS |
| ^1^ NS: Not significant | | | |

| Table S6: Variant Sequence Data and NCBI Results | | | | |
| --- | --- | --- | --- | --- |
| Variant | Sequence | Reported Hosts NCBI Blast | | |
| blaTEM  Variant 1 | >blaTEM_Variant_1  ATACCAAACGACGAGCGTGACACCACGATGCCTGTAGCAATGGCAACAACGTTGCGCAAACTATTAACTGGCGAACTACTTACTCTAGCTTCCCGGCAACAATTAATAGACTGGATGGAGGCGGATAAAGTTGCAGGACCACTTCTGCGCTCGGCCCTTCCGGCTGGCTGGTTTATTGCTGATAAATCTGGAGCCGGTGAGCGTGGGTCTCGCGGTATCATTGCAGCACTGGGG | *Achromobacter denitrificans*  *Acinetobacter baumannii*  *Acinetobacter haemolyticus*  *Acinetobacter sp.*  *Aeromonas hydrophila*  *Alcaligenes sp.*  *Achromobacter denitrificans*  *Acinetobacter baumannii*  *Acinetobacter haemolyticus*  *Acinetobacter sp.*  *Aeromonas hydrophila*  *Alcaligenes sp.*  *Arabidopsis thaliana*  *Babesia bigemina*  *Bacillus cereus*  *Bacillus mycoides*  *Bacillus subtilis*  *Bacillus safensis*  *Bacillus sp. BT-B158*  *Bacillus subtilis*  *Bacillus tropicus*  *Bacillus velezebsus*  *Bifidobacterium longum*  *Burkholderia cepacian*  *Burkholderia lata*  *Burkholderia sp.* | *Chlamydia trachomatis*  *Clostridioides difficile*  *Clostridium botulinum*  *Cronobacter sakazakii*  *Escherichia coli*  *Eimeria acervulina*  *Eimeria maxima*  *Enterobacter cloacae*  *Enterococcus faecium*  *Escherichia sp.*  *Escherichia marmotae*  *Faecalibacterium prausnitzii*  *Flavobacterium columnare*  *Francisella philomiragia*  *Helicobacter pylori*  *Klebsiella michiganensis*  *Klebsiella pneumoniae*  *Klebsiella quasipneumoniae*  *Legionella pneumophila*  *Macrococcus caseolyticus*  *Morganella morganii*  *Mycolicibacterium smegmatis*  *Mycobacterium tuberculosis*  *Mycoplasma mycoides*  *Neisseria meningitidis* | *Phaffia rhodozyma*  *Propionibacterium freudenreichii*  *Pseudomonas aeruginosa*  *Pseudomonas sp.*  *Rhizobium leguminosarum*  *Ruthenibacterium lactatiformans*  *Saccharomyces cerevisiae*  *Salmonella enterica*  *Schizosaccharomyces pombe*  *Serratia marcescens*  *Solanum pennellii*  *Staphylococcus aureus*  *Staphylococcus hominis*  *Staphylococcus saprophyticus*  *Streptococcus agalactiae*  *Streptococcus lutetiensis*  *Streptococcus pneumoniae*  *Cloning vectors*  *Expression vectors*  *Stenotrophomonas sp.*  *Streptomyces sp.*  *Trypanosoma brucei*  *uncultured bacterium*  *Vibrio parahaemolyticus*  *Vibrio vulnificus* |
| Variant | Sequence | Reported Hosts NCBI Blast | | |
| blaTEM  Variant 2 | >blaTEM Variant 2  ATACCAAACGACGAGCGTGACACCACGATGCCTGCAGCAATGGCAACAACGTTGCGCAAACTATTAACTGGCGAACTACTTACTCTAGCTTCCCGGCAACAATTAATAGACTGGATGGAGGCGGATAAAGTTGCAGGACCACTTCTGCGCTCGGCCCTTCCGGCTGGCTGGTTTATTGCTGATAAATCTGGAGCCGGTGAGCGTGGGTCTCGCGGTATCATTGCAGCACTGGGG | *Acinetobacter baumannii*  *Acinetobacter johnsonii*  *Acinetobacter towneri*  *Aeromonas caviae*  *Aeromonas hydrophila*  *Aeromonas media*  *Aeromonas sp. ASNIH2*  *Aeromonas veronii*  *Atlantibacter hermannii*  *Bacillus subtilis*  *Bacteroides fragilis*  *Chlamydia trachomatis*  *Chryseobacterium gallinarum*  *Citrobacter amalonaticus*  *Citrobacter braakii*  *Citrobacter farmeri*  *Citrobacter freundii*  *Citrobacter koseri*  *Citrobacter portucalensis*  *Citrobacter sp.*  *Citrobacter werkmanii*  *Citrobacter youngae*  *Clostridioides difficile*  *Cronobacter sakazakii*  *Enterobacter asburiae*  *Enterobacter bugandensis*  *Enterobacter chengduensis*  *Enterobacter cloacae*  *Enterobacter hormaechei* | *Enterobacter kobei*  *Enterobacter roggenkampii*  *Enterococcus faecium*  *Enterobacter sp.*  *Enterobacteriaceae bacterium*  *Escherichia albertii*  *Escherichia coli*  *Escherichia fergusonii*  *Escherichia marmotae*  *Haemophilus influenzae*  *Haemophilus parainfluenzae*  *Hafnia alvei*  *Kingella kingae*  *Klebsiella aerogenes*  *Klebsiella huaxiensis*  *Klebsiella michiganensis*  *Klebsiella oxytoca*  *Klebsiella pneumoniae*  *Klebsiella quasipneumoniae*  *Klebsiella sp.*  *Klebsiella variicola*  *Leclercia adecarboxylata*  *Leclercia adecarboxylata*  *Leclercia sp.*  *Morganella morganii* | *Mycobacterium tuberculosis*  *Neisseria gonorrhoeae*  *Neisseria mucosa*  *Proteus mirabilis*  *Proteus terrae subsp. cibarius*  *Proteus vulgaris*  *Providencia huaxiensis*  *Providencia rettgeri*  *Pseudomonas aeruginosa*  *Pseudomonas fluorescens*  *Raoultella ornithinolytica*  *Raoultella planticola*  *Salmonella enterica*  *Salmonella sp.*  *Serratia liquefaciens*  *Serratia marcescens*  *Serratia sp.*  *Shigella boydii*  *Shigella flexneri*  *Shigella sonnei*  *Shigella sp.*  *Staphylococcus aureus*  *Streptococcus pneumoniae*  *Synthetic construct*  *Uncultured bacterium*  *Vibrio cholerae*  *Vibrio parahaemolyticus*  *Vibrio vulnificus*  *Yersinia pseudotuberculosis* |
| Variant | Sequence | Reported Hosts NCBI Blast | | |
| blaTEM  Variant 3 | >blaTEM Variant 3  ATACCAAACGACGAGCGTGACACCACGATGCCTGCAGCAATGGCAACAACGTTGCGCAAACTATTAACTGGCGAACTACTTACTCTAGCTTCCCGGCAACAATTAATAGACTGGATGGAGGCGGATAAAGTTGCAGGACCACTTCTGCGCTCGGCCCTTCCGGCTGGCTGGTTTATTGCTGATAAATCTGGTGCCGGTGAGCGTGGGTCTCGCGGTATCATTGCAGCACTGGGG | Synthetic construct |  |  |
| pmrC  Variant1 | >pmrC Variant 1  TTGATGTACTCTTCAAGCCCGTGGAACAGCACTTCGTCATAGCATTCGCCGTTGATGCACTGATCAGGTAGATTCAGCGCGGTGACGTTCTGGTGAGGCACGCGGTCGCAGGCACCTTTACAGCCGCCATCGTTGTCATTCCACAGCACGTTGATGCCCGCTCGCTGAATGATATCCAGCACGCCTTCCTGGTGCTGTGCCAGCTCTTCTTTGTAGTGCTCACGCGGCATATCCGAGAA | *Escherichia coli*  *Escherichia sp.*  *Salmonella sp.*  *Shigella boydii*  *Shigella flexneri*  *Shigella sonnei*  *Synthetic construct* |  |  |
| pmrC  Variant2 | >pmrC Variant 2  TTGATGTACTCTTCCAGCCCGTGGAACAGTACTTCGTCATAGCATTCGCCGTTGATGCACTGACCAGGCAGATTCAGCGCGGTGACGTTCTGGTGAGGCACGCGGTCGCAGGCACCTTTACAGCCGCCATCGTTGTCATTCCACAGCACGTTGATGCCCGCTCGCTGAATGATATCCAGCACGCCTTCCTGGTGCTGTGCCAGCTCTTCTTTGTAATGCTCACGCGGCATATCCGAGAA | *Escherichia coli*  *Salmonella sp.*  *Shigella flexneri*  *Shigella sonnei*  *Shigella sp.* |  |  |
| Variant | Sequence | Reported Hosts NCBI Blast | | |
| pmrC  Variant 3 | >pmrC Variant 3  AAACGACGGATTGTTGTAATGGTTGCCAATACGCGGATCGAGACAAACGCCTTCGATAATCTGGCGAGTATTCAGACCTAAACTTTCTGCATAGCTATCCAGTTCATTAAAGTACGCTACGCGCATCGCCAGATAGGTATTAGCGAAAAGTTTTATCGCTTCTGCTTCAGTGGAGTCGGTAAACAGGGTCGGGATATTTTGCTTAATCGCCCCTTCCTGTAACAACGCAGCAAAACGTTCAGCGC | *Escherichia coli* |  |  |
| pmrC  Variant 4 | >pmrC Variant 4  TTGATGTACTCTTCCAGCCCGTGGAACAGCACTTCGTCATAGCATTCGCCGTTGATGCACTGACCAGGCAGGTTCAGCGCGGTGACGTTCTGGTGAGGTACGCGATCGCAAACGCCTTTACAGCCGCCATCGTTGTCTTTCCACAGCACGTTGATGCCCGCTCGCTGAATGATATCCAGCACGCCTTCCTGGTGCTGTGCCAGCTCTTCTTTGTAGTGCTCACGCGGCATATCCGAGAA | *Escherichia coli* |  |  |
| blaOXA  Variant 1 | >blaOXA Variant 1  GCAATGGGAAAGAGACTTGACCTTAAGAGGGGCAATACAAGTTTCAGCTGTTCCCGTATTTCAACAAATCGCCAGAGAAGTTGGCGAAGTAAGAATGCAGAAATACCTTAAAAAATTTTCCTATGGCAACCAGAATATCAGTGGTGGCATTGACAAATTCTGGTTGGAAGGCCAGCTTAGAATTTCCGCAGTTAATCAAGTGGAGTTTCTAGAGTCTCTATATTTAAATAAATTGTCAGCATCTAAAGAAAACCAGCTAATAGTAAAAGAGGCTTTGGTAACGGAGGCGGCACCTGAATATCTAGTGCATTCAAAAACTGGTTTTTCTGGTGTGGGAACTGAG | *Acinetobacter baumannii*  *Acinetobacter johnsonii*  *Aeromonas caviae*  *Aeromonas hydrophila*  *Aeromonas media*  *Aeromonas salmonicida*  *Aeromonas simiae*  *Aeromonas sp.*  *Aeromonas veronii*  *Citrobacter braakii*  *Citrobacter freundii*  *Citrobacter sedlakii*  *Citrobacter werkmanii*  *Enterobacter aerogenes*  *Enterobacter asburiae*  *Enterobacter cloacae*  *Enterobacter hormaechei*  *Enterobacter kobei* | *Escherichia coli*  *Escherichia fergusonii*  *Gallibacterium anatis*  *Klebsiella grimontii*  *Klebsiella michiganensis*  *Klebsiella oxytoca*  *Klebsiella pneumoniae*  *Klebsiella quasipneumoniae*  *Morganella morganii*  *Pantoea agglomerans*  *Proteus mirabilis*  *Proteus sp.*  *Proteus vulgaris*  *Providencia heimbachae*  *Providencia rettgeri*  *Providencia sp.*  *Providencia stuartii*  *Pseudocitrobacter faecalis* | *Pseudomonas aeruginosa*  *Pseudomonas fulva*  *Pseudomonas monteilii*  *Pseudomonas putida*  *Pseudomonas shirazica*  *Pseudomonas sp.*  *Salmonella enterica*  *Salmonella sp.*  *Serratia marcescens*  *Shewanella putrefaciens*  *Shigella flexneri*  *Shigella sonnei*  *Stenotrophomonas maltophilia*  *Vibrio cholerae*  *Vibrio fluvialis*  *Vibrio parahaemolyticus*  *Yokenella regensburgei* |
| Variant | Sequence | Reported Hosts NCBI Blast | | |
| blaOXA_Variant2 | >blaOXA Variant 2  ACAATGGGAAAGAGACTTGAGCTTAAGAGGGGCAATACAAGTTTCAGCGGTTCCCGTATTTCAACAAATCGCCAGAGAAGTTGGCGAAGTAAGAATGCAGAAATATCTTAAAAAATTTTCATATGGTAACCAGAATATCAGTGGTGGCATTGACAAATTCTGGTTGGAGGGTCAGCTTAGAATTTCCGCAGTTAATCAAGTGGAGTTTCTAGAGTCTCTATTTTTAAATAAATTGTCAGCATCAAAAGAAAATCAGCTAATAGTAAAAGAGGCTTTGGTAACGGAGGCTGCGCCTGAATATCTTGTGCATTCAAAAACTGGTTTTTCTGGTGTGGGAACTGAG | *Aeromonas caviae*  *Aeromonas hydrophila*  *Alcaligenes faecalis*  *Citrobacter freundii*  *Klebsiella oxytoca*  *Klebsiella michiganensis*  *Proteus mirabilis*  *Pseudomonas aeruginosa* |  |  |
| blaOXA_Variant3 | >blaOXA Variant 3  GAACGCCAAGCGGATCGTGCCATGTTGGTTTTTGATCCTGTGCGATCGAAGAAACGCTACTCGCCTGCATCGACATTCAAGATACCTCATACACTTTTTGCACTTGATGCAGGCGCTGTTCGTGATGAGTTCCAGATTTTTCGATGGGACGGCGTTAACAGGGGCTTTGCAGGCCACAATCAAGACCAAGATTTGCGATC | *Achromobacter denitrificans*  *Acientobacter baumannii*  *Aeromonas caviae*  *Aeromonas sp.*  *Alcaligenes faecalis*  *Bordetella bronchiseptica*  *Citrobacter freundii*  *Corynebacterium asperum*  *Enterobacter cloacae*  *Enterobacter hormaechei*  *Enterobacter kobei* | *Escherichia coli*  *Enterobacter aerogenes*  *Klebsiella oxytoca*  *Klebsiella pneumoniae*  *Klebsiella quasipneumoniae*  *Klebsiella sp.*  *Mannheimia haemolytica*  *Morganella morganii*  *Pasteurella multocida*  *Phytobacter ursingii*  *Providencia rettgeri* | *Pseudomonas aeruginosa*  *Pseudomonas baetica*  *Pseudomonas monteilii*  *Pseudomonas putida*  *Salmonella enterica*  *Salmonella typhimurium*  *Serratia marcescens*  *Staphylococcus aureus*  *Stenotrophomonas maltophilia*  *Uncultured bacterium*  *Vibrio anguillarum*  *Vibrio cholerae* |
| blaOXA_Variant4 | >blaOXA Variant 4  GAACGCCAAGCGGATCGTGCCATGTTGGTTTTTGATCCTGTGCGATCGAAGAAACGCTACTCGCCTGCATCGACATTCAAGATACCTCATACACTTTTTGCACTTGATGCAGGCGCTGTTCGTGATGAGTTCCAGATTTTTCGATGGGACGGCGTTAACAGGGGCTTTGCAGGCCACAATCAAGACCAAGATTTT | No 100% similarity observed |  |  |
| Variant | Sequence | Reported Hosts NCBI Blast | | |
| mdtH  Variant1 | >mdtH Variant 1  TTGAAGCGTGTCTGTCGTTAACGTTGCTCTACCCTATCGCCCGCTGGAGTGAAAAGCATTTTCGTCTGGAACACCGGTTG  ATGGCTGGGCTGTTGATAATGTCATTAAGCATGATGCCGGTGGGCATGGTCAGCGGCCTGCAACAACTTTTCACCCTGAT  TTGTCTGTTTTATATCGGGTCGATCATTGCCGAGCCTGCGCGTGAAACCTTAAGTGCTTCGCTGGCGGACGCAAGAGCTC  GCGGCAGCTATATGGGGTTTAGC | *Escherichia coli*  *Escherichia fergusonii*  *Escherichia sp.*  *Shigella boydii*  *Shigella dysenteriae*  *Shigella felxneri*  *Shigella sonnei*  *Synthetic Escherichia coli* |  |  |
| mdtH  Variant2 | >mdtH Variant 2  TTGAAGCGTGTCTGTCGTTAACGTTGCTCTATCCTATCGCCCGCTGGAGTGAAAAGCATTTTCGTCTGGAACACCGGTTGATGGCTGGGCTGTTGATAATGTCATTAAGCATGATGCCGGTGGGCATGGTCAGCGGCCTGCAACAACTTTTCACCCTGATTTGTCTGTTTTATATCGGGTCGATCATTGCCGAGCCTGCGCGTGAAACCTTAAGTGCTTCGCTGGCGGACGCAAGAGCTCGCGGCAGCTATATGGGGTTTAGC | *Escherichia coli*  *Shigella boydii*  *Shigella flexneri*  *Shigella sp.* |  |  |
| mdtH  Variant3 | >mdtH Variant 3  TTGAAGCGTGTCTGTCGTTAACGTTGCTCTACCCTATCGCCCGCTGGAGTGAAAAGCATTTTCGTCTGGAACACCGGTTAATGGCTGGGCTGTTGATAATGTCATTAAGCATGATGCCAGTAGGCATGGTCAGCGGCCTGCAACAACTTTTCACCCTGATTTGTCTGTTTTATATCGGGTCGATCATTGCCGAGCCTGCGCGTGAAACCTTAAGTGCTTCGCTGGCAGACGCAAGAGCTCGCGGCAGCTATATGGGGTTTAGC | *Enterobacter hormaechei*  *Escherichia coli*  *Shigella flexneri* |  |  |
| Variant | Sequence | Reported Hosts NCBI Blast | | |
| mdtH  Variant4 | >mdtH Variant 4  TTGAAGCGTGTCTGTCGTTAACGTTGCTCTATCCTATCGCCCGCTGGAGTGAAAAGCATTTTCGTCTGGAACATCGGTTGATGGCTGGGCTGTTGATAATGTCATTAAGCATGATGCCAGTGGGCATGGTCAGCGGCCTGCAACAACTTTTCACCCTGATTTGTCTGTTTTATATCGGGTCGATCATTGCCGAGCCTGCGCGTGAAACCTTAAGTGCTTCGCTGGCAGACGCAAGAGCTCGCGGCAGCTATATGGGGTTTAGC | *Escherichia coli*  *Shigella dysenteriae* |  |  |
